# General Features of Transmembrane Beta Barrels From a Large Database

**DOI:** 10.1101/2022.04.13.488089

**Authors:** Daniel Montezano, Rebecca Bernstein, Matthew M. Copeland, Joanna S. G. Slusky

**Author notes:** University of California, Berkeley, CA 94720. **Author Contributions:** J.S.G.S. and D.M. designed the research; D.M., and R.B. performed the research, analyzed the data and wrote the paper; M.M.C. contributed software tools. J.S.G.S and D.M. edited the paper.

## Abstract

Large datasets contribute new insights to subjects formerly investigated by exemplars. We used co-evolution data to create a large, high-quality database of transmembrane β-barrels (TMBB). By applying simple feature detection on generated evolutionary contact maps, our method (IsItABarrel) achieves 95.88% balanced accuracy when discriminating among protein classes. Moreover, comparison with IsItABarrel revealed a high rate of false positives in previous TMBB algorithms. In addition to being more accurate than previous datasets, our database (available online) contains 1,894,206 bacterial TMBB proteins from 20 evolutionary classes, respectively 17 and 2.2 times larger than the previous sets TMBB-DB and OMPdb. We anticipate that due to its quality and size the database will serve as a useful resource where high quality TMBB sequence data is required. We found that TMBBs can be divided into 10 types, three of which have not been previously reported. We find tremendous variance in proteome percentage among TMBB-containing organisms with some using 6.79% of their proteome for TMBBs and others using as little as 0.27% of their proteome. The distribution of the lengths of the TMBBs is suggestive of previously hypothesized duplication events. In addition, we find that the C-terminal β-signal varies among different classes of bacteria though it is most commonly HyGHyGY+F. However, this β-signal is only characteristic of prototypical TMBBs. The nine non-prototypical barrel types have other C-terminal motifs and it remains to be determined if these alternative motifs facilitate TMBB insertion or perform any other signaling function.

**Significance Statement:** Outer membrane proteins (OMPs) control all interactions between Gram negative bacteria and their environments including uptake and efflux of antibiotics. We created an algorithm that identifies bacterial OMPs from sequence. The quality of our algorithm allows us to identify most OMPs (∼1.8 million) from prokaryotic genomes including >240,000 unrelated to previously structurally-resolved OMPs. We identify ten types of OMPs in our database. The largest type’s signal sequence—used for targeting the membrane-insertion machinery—varies by phylogenetic class. All other types of OMPs do not have a related signal sequence, raising new questions of how these proteins fold. Our web-accessible database will allow for further exploration of the varieties of outer membrane proteins to uncover new targets for controlling antibiotic resistance.

## Introduction

Comparative studies of organisms, genomes, and protein families are dependent on the quality and size of biological databases. These studies form the basis for answering questions about the evolutionary history of different protein families and for identifying and classifying the diversity of protein structure and function. The proteins found in the outer membrane of Gram-negative bacteria, generally have a highly similar β-barrel fold (1), which has evolved mostly by replication of an ancestral hairpin motif. (2, 3) Despite the significant structural similarity, this fold emerges from a rich diversity of primary sequences (4). Attempts to classify the barrel fold into families has met with a number of challenges that can be observed in the diversity of current classifications performed in public structural databases such as SCOPe. The functional roles of transmembrane β-barrels (TMBBs) are also diverse, including nutrient uptake, membrane stabilization, catalysis, cell adhesion, cell signaling, and efflux. (5)

The hydrophobic environment of the outer membrane imposes constraints on the sequence and topology of TMBB proteins. Since the outside of barrels is in contact with the membrane and the inside of the barrel with the water-filled lumen, strand residues that point into the channel are usually hydrophilic, while residue side chains pointing outside into the membrane are hydrophobic and aromatic. Such constraints induce an imperfect hydrophobicity alternation pattern on residues of each strand. (6) Also, there are significant differences in amino acid abundance at different depths across the membrane, with more hydrophobic residues in the central region of the membrane. (7-9) Attempts have been made to computationally identify TMBBs using amino acid composition (10-12), using specific knowledge about TMBBs and their environment (7, 13, 14) and by identification of sequence motifs. (15, 16) In addition to classification within the fold, several computational methodologies also predict more specific features of TMBB topology such as strand number and which parts of the protein localize to the periplasm, to the outside of the cell in the extracellular milieu or are embedded in the membrane. (14, 17-21)

Homology has been explored and combined with many of these previous methods to augment available data and make explicit use of evolutionary cues in order to improve TMBB classification and prediction. (22-24) Machine learning methods were also employed to the tasks of TMBB topology prediction and fold classification, including feed-forward neural networks and SVMs (25), recurrent neural networks (26), hidden Markov models (27, 28), extreme learning machines (29) and ensemble approaches. (26, 30)

In two instances algorithms were applied to establish large-scale homologous databases of TMBBs or to perform proteome-wide search for new TMBBs. The TMBB-DB (31) contains 1,881,712 sequences collected from 600 different bacterial proteomes with sequences ranked by their likelihood of encoding a TMBB. The OMPdb (32) contains 1,198,558 (as of Jan 2021) protein sequences predicted to be TMBBs by the PRED-TMBB2 algorithm.

These previous methods were dependent on the availability of experimentally solved structures, which is scarce for β-barrels. Methods that used sequences were trained on a comparatively small set of putative TMBB sequences from which sequence features were extracted. Overall, these methods used protein information in a limited sense.

Co-evolution information is obtained from analyzing covariation of residues in an alignment of protein sequences that are homologous to a protein of interest. Strong statistical co-variation of two or more residues indicates a possible co-evolutionary event, where one set of mutations is compensated by another set of mutations. (33) A co-evolutionary event can be interpreted as evidence that the residues are in proximity, from which contact probabilities can be derived (34). If higher order interactions can be reliably extracted, contacts can be predicted with higher accuracy. (35) Evolutionary contact maps are 2D representations that show the likelihood of two residues in a protein sequence being in contact and may be used for predicting protein structure or for refining homology models. Here we use contact information to discriminate between the barrel fold and other protein folds.

Although contact information can be extracted from protein structure models, generated for example, by AlphaFold (36), this solution still does not provide a reliable way to categorize the predicted structures as barrels. We would still require an algorithm that could analyze these predicted structures in detail and decide if they represent a β-barrel structure or not. Currently, there is not yet a comprehensive categorization of AlphaFold structures into SCOPe families. Moreover, SCOPe scatters TMBBs among a variety of different folds (4), making it difficult to track down all instances of the fold.

Here we present a new methodology called IsItABarrel that uses co-evolution information and extracts discriminative features from contact maps for the identification of TMBBs from sequences. We apply IsItABarrel to identify 1,894,206 TMBB proteins from 14,056 bacterial organisms. In addition, we used IsItABarrel to search 2,959 bacterial proteomes for TMBBs and provide updated estimates of the TMBB content in these organisms. We also analyze general features of our new, large TMBB dataset and find evolutionary class-specific β-signals. Finally, we find ten different types of TMBBs each with their own length distribution, organism preference, and β-signal. Three of the ten TMBB types have not been previously documented.

## Results

### Algorithm for TMBB Prediction

Up-down β-barrels such as those found in bacterial outer membranes have contact maps that can be distinguished from other proteins (Figs. 1 A and B). The two features that nearly definitively identify TMBBs from contact maps are strand-strand interactions (Fig. 1C) and closing contacts (Fig. 1D). Our algorithm, called IsItABarrel, identifies TMBBs by either the existence of a closing contact or the existence of at least four β-strand contacts of 16-25 contact map residues each (Fig. 1E and Fig. S1). We use the Matthew’s correlation coefficient (MCC) (37) to measure algorithm performance because it is less skewed by imbalanced datasets. We validated IsItABarrel on a set of 1,121 proteins with solved crystal structures, 121 of which are TMBBs and 1000 are non-TMBBs, yielding 0.921 MCC (95.88% balanced accuracy) with eight instances of false positives and nine instances of false negatives (Table 1, left). In addition, we validated IsItABarrel on a dataset containing only signal-sequence containing proteins and measured an MCC of 0.778 (balanced accuracy of 94.91%) (Table 1, right). We compared IsItABarrel with other predictors used to create previous datasets: the Freeman-Wimley algorithm that was used to create the TMBB-DB database (13), PRED-TMBB2 (20) that was used to create the OMPdb (32) and also with the BOMP predictor (15). We find that IsItABarrel has the highest MCC and the fewest false positives (Table 1 and Fig. S2), thereby allowing it to generate the most reliable dataset.

**Figure 1.**
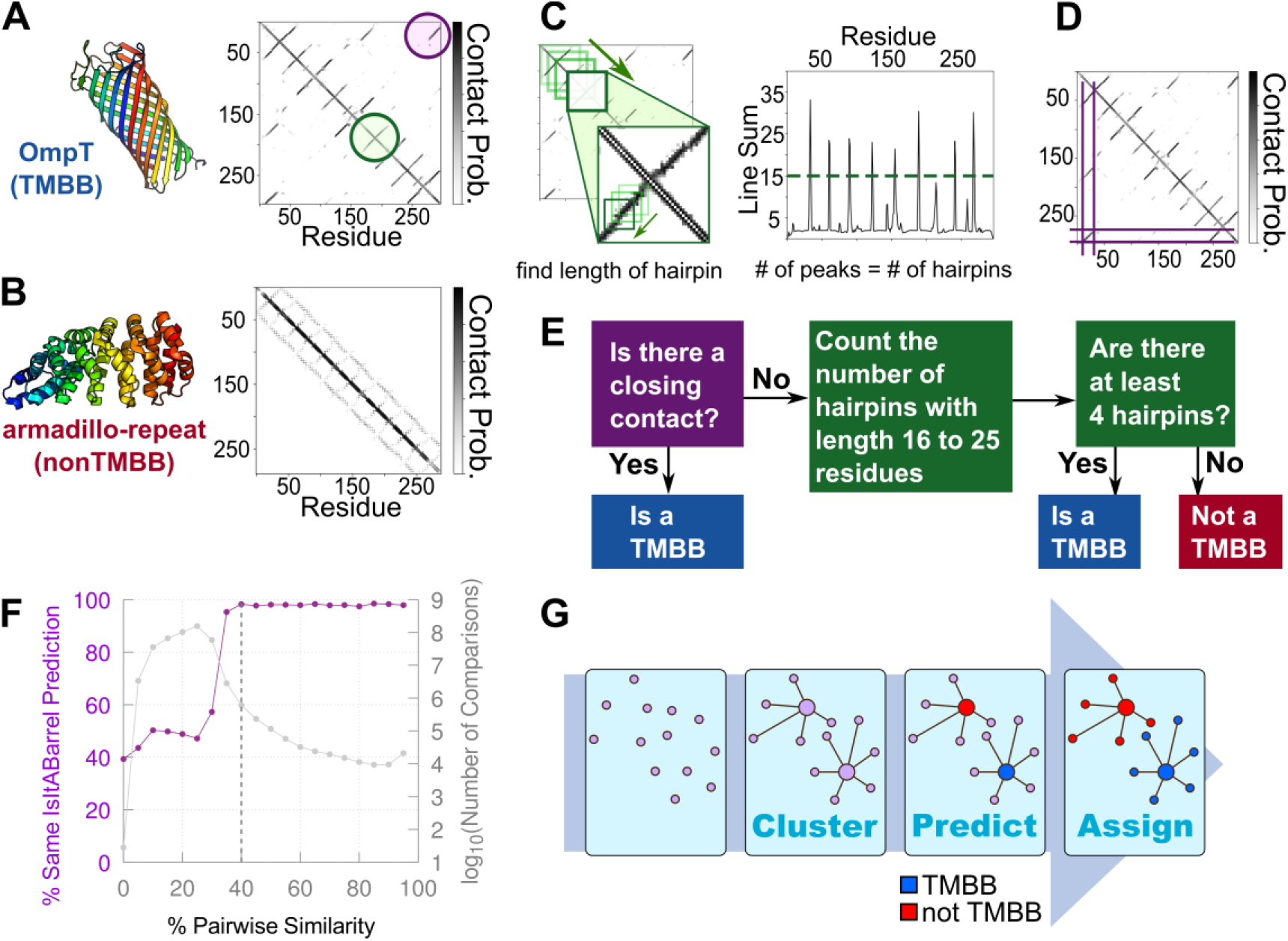
The IsItABarrel Algorithm. **(A and B)** Structures are colored from N to C-terminus using rainbows from blue to red. Probability of contact is indicated by pixel grey value, with darker pixel indicating stronger contact between residues in *x* and *y* axes. **(A)** OmpT (PDB ID: 1I78) (left) and corresponding contact map predicted by RaptorX-Contact (right). The map shows typical features of β-barrel proteins, anti-parallel contacts between neighboring strands (green circle) and the closing contact between the first and last strand of the barrel (purple circle). **(B)** Structure of the non-TMBB armadillo-repeat protein (PDB ID: 4V3O) (left) and corresponding contact map predicted by RaptorX-Contact (right) without the typical features found in maps of β-barrels. **(C)** Strand-strand interaction (β-hairpin) detection. Map is scanned along the main diagonal in search of β-hairpins. Detected β-hairpins are characterized in terms of the length of their contact network (inset). The sum of the contact probabilities along a successfully detected β-hairpin is compared against the threshold for inclusion in the final TMBB classification step. **(D)** A closing contacts line is detected that aligns with the first and last strands **(E)** Classification by IsItABarrel is a two-step process requiring either a closing contact or four hairpins. **(F)** Sequences sharing more than 40% identity tend to have the same IsItABarrel classification. Percentage of pairs that share the same IsItABarrel prediction (y-axis) for each bin of percent similarity (x-axis) (purple), total number of comparisons (log_10_) at each similarity level (gray). Dashed line at 40% pairwise sequence similarity. **(G)** Steps the algorithm used to assign IsItABarrel predictions by similarity. A large set of sequences is clustered by sequence similarity. IsItABarrel predictions are made for the cluster representatives. Cluster members are assigned the same prediction as their cluster representative.

**Table 1.** Comparison of predictors on validation sets.

|  | Dataset |  |  |  |  |  |  |  |  |  |
| --- | --- | --- | --- | --- | --- | --- | --- | --- | --- | --- |
|  | NONTMBB1K<br>(121 TMBBs / 1000 non-TMBBs) |  |  |  |  | SS1929<br>(121 TMBBs / 1929 non-TMBBs) |  |  |  |  |
| Algorithm | MCC | TP | TN | FP | FN | MCC | TP | TN | FP | FN |
| IsItABarrel | 0.921 | 112 | 992 | 8 | 9 | 0.778 | 112 | 1876 | 53 | 9 |
| BOMP | 0.863 | 111 | 979 | 21 | 10 | 0.647 | 111 | 1815 | 114 | 10 |
| FW-BB<br>Analysis | 0.756 | 121 | 925 | 75 | 0 | 0.460 | 121 | 1582 | 347 | 0 |
| PRED-TMBB2 | 0.791 | 117 | 947 | 53 | 4 | 0.588 | 117 | 1752 | 177 | 4 |

Left column, NONTMBB1K, includes 121 non-homologous TMBBs as positive examples and 1,000 non-homologous non-TMBBs as the negative set. Right column, SS1929, includes the same set of 121 TMBBs and a different set with only non-TMBB membrane proteins, identified with SignalP5.0. IsItABarrel (top row) provides the lowest number of false positives and intermediate performance for true positives. TP – true positives; TN – true negatives; FP – false positives; FN – false negatives. Matthews' Correlation Coefficient (MCC) calculated as: $MCC = (TP \cdot TN - FP \cdot FN) / \sqrt{((TP + FP)(TP + FN)(TN + FP)(TN + FN))}$ .

### Prediction of a Large Dataset with IsItABarrel

Map creation is a computationally demanding process. To increase the prediction scale, we determined the sequence similarity level that maintained consistent IsItABarrel classification. We determined that new maps do not need to be made for proteins with 40% or greater pairwise sequence similarity as homologs above that threshold have above a 95% similar IsItABarrel prediction (Fig. 1F), a success rate similar to the balanced accuracy of our algorithm overall. We determined this metric using the TMBB29183 dataset (Table S1) by evaluating 425,809,153 pairwise sequence comparisons and obtaining a distribution of sequence similarity values ranging from 4% to 99.9%. Similar IsItABarrel evaluations for proteins with high sequence similarity also indicate that our algorithm is robust for identifying bacterial TMBBs as more related proteins are likely to have the same fold.

### Expanding the Dataset

By combining and evaluating the proteins in previous datasets (OMP-DB and TMBB-DB) we created a database of 514,728 bacterial TMBB proteins. We then undertook a more thorough search of genetic space to uncover previously unannotated bacterial TMBBs.

We considered 2,959 NCBI prokaryote reference and representative organisms. These proteomes span 78 classes and 35 phyla (Tables S2 and S3). Around 400 of these organisms were not diderms and did not have any TMBBs. We found consistent representation of TMBBs (*i*.*e*. TMBBs in ten or more organisms) in 20 classes. Six of the classes were in two phyla (Bacteroidetes and Firmicutes) and within those phyla they had consistent enough representation of TMBBs that we could represent them as phyla rather than individual classes. Thus our analysis comprises 2 phyla and 14 classes (Fig. 2). TMBB-containing organismal categories included those that are known to include monoderms (Actinomycetia and Firmicutes). Other classes, such as Thermotogae and Deinococci, are diderms but usually contain atypical membrane compositions or thicker peptidoglycan layers which set them apart from other diderms. (38)

**Figure 2.**
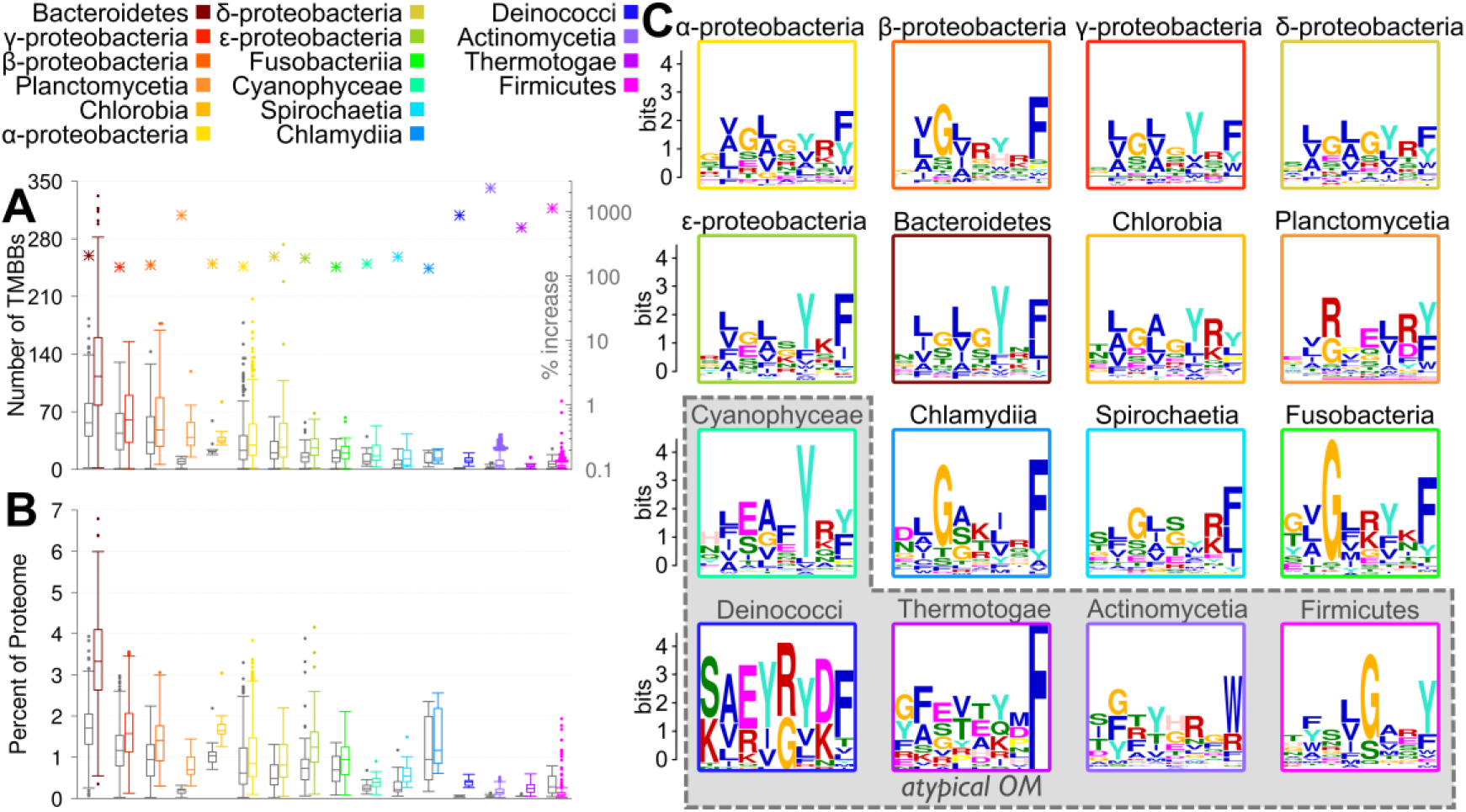
TMBB Content in Bacterial Proteomes and Predicted C-terminal β-signal Motif. **(A and B)** Colored by class/phylum, categories are sorted by decreasing median of number of TMBBs. **(A)** Distribution of number of TMBBs in 2,556 reference and representative bacterial proteomes. TMBBs predicted with IsItABarrel that were within other datasets (gray), and updated numbers after a full search with IsItABarrel (color). Number of TMBB proteins per organism by class/phylum (boxplot), percent increase in number of TMBBs from the initial search (stars using log_10_ scale at right). **(B)** Distribution of percentage of proteome predicted to encode TMBB proteins (same organisms as in A). **(C)** Logos plots of motifs found in the C-terminus of TMBBs from each phyla/class. The letters at each position are ordered from top to bottom by most common to least common. The height of each letter is proportional to its frequency in the column, while the height of the stack is adjusted to be the information content expressed in bits of the aligned sequences at that particular position. The colors of the letters indicate physicochemical properties: orange, small, magenta, acidic, blue, hydrophobic plus cysteine, cyan, tyrosine, pink, histidine, green, polar, red, basic, yellow, proline. Terrabacteria with gray background, Gracilicutes with white background.

We searched those TMBB-containing proteomes for possible new TMBBs by extracting all membrane proteins from the largest organism (in terms of proteome size) in each class/phylum. We predicted the presence of a signal peptide and kept in our set only the proteins that were predicted to have a signal peptide. We then predicted all of the remaining sequences of putative membrane proteins with IsItABarrel. In this search a total of 43,822 new protein sequences were found to be potential TMBBs, producing a 71.3% increase over what we had previously found (Figs. 2 A and B). To find the homologs of these new-found TMBBs, we performed a new search starting with the non-redundant protein database from NCBI. This search resulted in our final set of predicted TMBBs we call IsItABarrelDB, containing 1,894,206 bacterial TMBB sequences, out of the total set of known proteins (∼230 million sequences).

### Distribution of TMBBs in Representative Organisms

A highly accurate dataset of TMBBs allows for the quantification of the TMBB distribution among different organisms (Fig. 2 A and B). We find that Bacteroidetes tend to have the most TMBBs with an average of 127.5 per organism (average of 3.4% of the proteome). Thermotogae, which is known to have an atypical outer membrane (38), has the fewest with an average of 5.28 per organism (average of 0.28% of the proteome), though we find 5.68-times more barrels in this class than previously identified.

### β-signal by Organism

The C-termini of TMBBs are known to have a sequence motif (39) that assists membrane insertion by interacting with the outer membrane translocation protein BAM (40-42) called a β-signal. (43) We analyzed the predicted TMBB sequences for the presence of the β-signal in the reference genomes using the MEME (44) sequence-motif detector (Fig. 2C). MEME searched the final quarter of each sequence so that the signal could be detected even if it is not in the terminal strand. (45) Though there are differences among the sequence motifs, many features are consistent. Eleven of fifteen classes/phyla most commonly have a phenylalanine at the C-terminal position (-1) of the motif. The other four have a tyrosine. Even more likely (13 out of 16 classes/phyla) is a charged amino acid at the penultimate position (-2). The other even positions (-4, -6, and -8) tend to have small amino acids like glycine in Gracilicutes and charged amino acids in Terrabacteria. Tyrosine is the most common amino acid in the -3 position for 12 of 16 classes/phyla. The overall β-signal sequence is H_y_GH_y_GY+F (where H_y_ is hydrophobic amino acids L or V, and + is positively charged amino acids K or R). Chlamydiia has an additional small amino acid position at -5, consistent with previously reported Fim/Usher β-signals. However, the Actinomycetia class has none of these features and has a β-signal more consistent with the previously identified LamB-like β-signal. (42)

### Clustering of Sequences in IsItABarrelDB to Known TMBBs and Other Types

We performed an analysis of sequence features on the IsItABarrelDB including a C-terminal motif MEME analysis as shown for the 16 taxonomic groups in Fig. 2. We aimed to identify groupings of proteins related to each other which we will call “types” to disambiguate from other phylogenetic groupings. We identified related proteins to structurally resolved-TMBBs with an iterative sequence similarity search. Types were merged when one or more sequences matched in both TMBB types (described in Supplemental Information, *Analysis of Sequences in IsItABarrelDB*). All proteins that could not be clustered with previously known types were merged. These proteins that were unrelated to known barrels were then clustered and the largest groups containing more than 3,000 proteins were selected for downstream analysis.

Through this process we found ten types of barrels, five that clustered with structurally-resolved TMBBs (*Prototypicals, Fim/Usher, LptD-like, LpxR-like* and *Tsx-like)*, and five types that did not cluster with structurally-resolved TMBBs. Two of these non-structurally-resolved types had functional associations in NCBI (*NfrA-like* and *PorT-like*) and three TMBB types were either associated with a domain of unknown function (DUF) or were annotated as hypothetical proteins (*DUF1302, Hypothetical Protein Type 1*, and *Hypothetical Protein Type 2)*.

The length distribution of the protein types (Fig. 3A) reveals differences among the types as well as evolutionary patterns. In the Prototypical type there is a three-mode distribution each approximately double the length of the previous. This is possibly indicative of the 4-stranded (oligomer) to 8-stranded to 16-stranded doubling events of TMBBs described previously. (3, 4) Bimodality is also present in the NfrA-like protein type. The Fim/Usher proteins have a non-doubling bimodal distribution, while most other types are unimodal (Fig. 3A).

**Figure 3.**
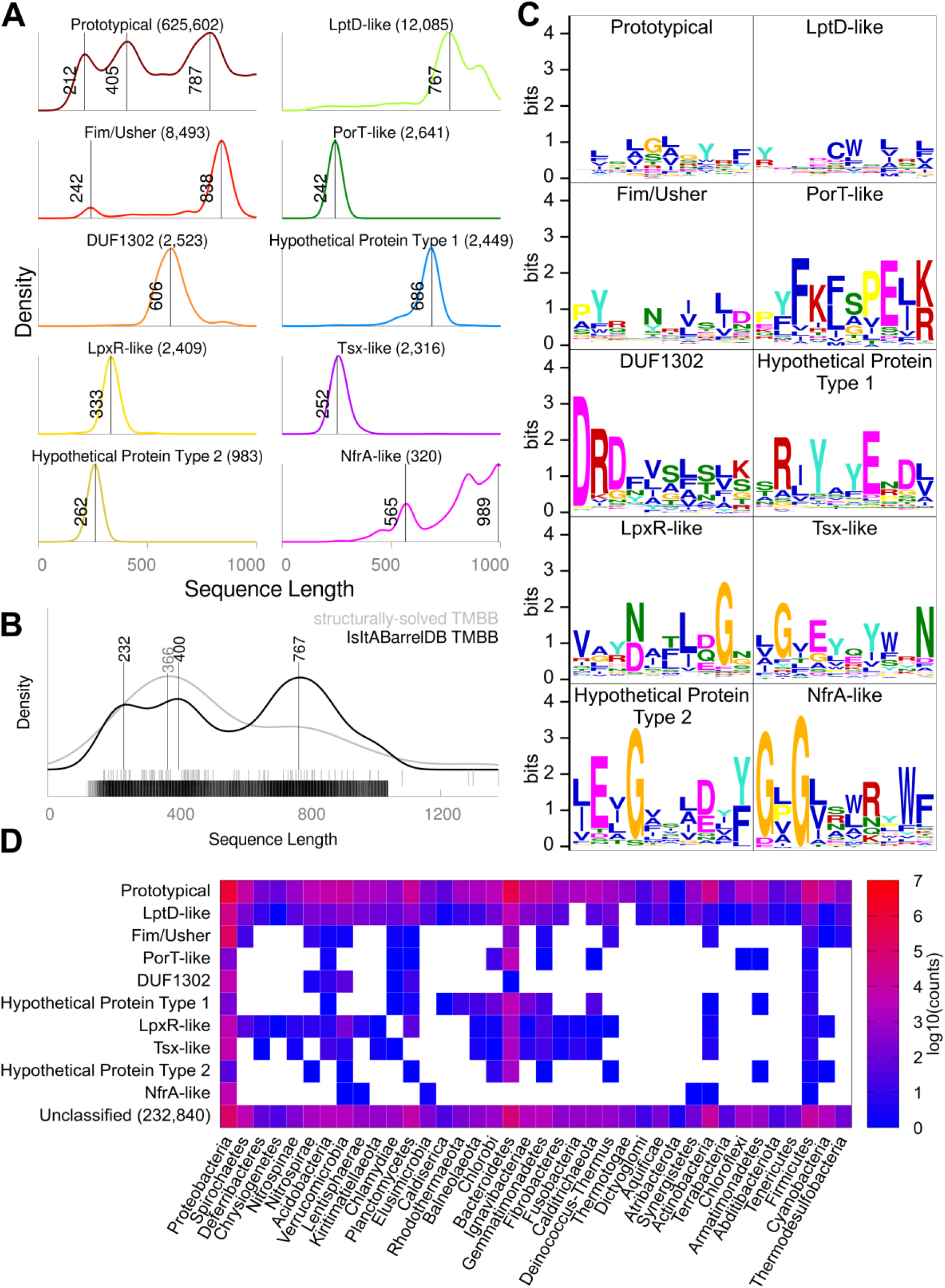
Analysis of IsItABarrelDB. **(A)** Distribution of sequence lengths of ten largest types of TMBBs in the IsItABarrelDB. **(B)** Kernel density estimate of the distribution of sequence lengths for structurally-solved (gray) and IsItABarrel predicted (black) TMBBs, rug plot showing instances below. **(C)** C-terminal sequence motifs found by MEME. Motif search was performed on a small subset of non-homologous representatives after clustering down to 40% sequence similarity (set sizes in Table S7). Sequence logos as described in Fig. 2C. **(D)** The protein sequences in the IsItABarrelDB are found in 38 phyla, with larger numbers in Proteobacteria and Bacteroidetes. While Prototypicals and LptD-like are found in all phyla, the other protein types are restricted to certain lineages. The Unclassified type across all phyla includes 232,840 sequences.

When comparing the IsItABarrelDB to 100 non-homologous structurally-resolved TMBBs we find that the most frequent size of TMBBs is 767 amino acids with another two modes for smaller barrels at 232 and 400 residues (Fig. 3B). The main peak of structurally solved proteins is at 366. Thus, while the smaller sized TMBBs are well represented in the structurally resolved set (peak of distribution at 366 amino acids) the larger barrels are not. In addition, though the structurally-resolved set has one mode, the full IsItABarrelDB has three modes. Notably, these modes are all roughly double each other, again suggestive of two possible duplications, though only the 8-stranded barrel to 16-stranded barrel duplication has been seen previously. (3) Since our analysis only includes 120 – 1040 residues we are likely missing a population of larger TMBBs that would be less tractable to evolutionary contact map analysis.

We find prototypical motifs dominated the sequence motifs in the β-signal class analysis. However, different types harbor different sequence motifs at their C-terminal end (Fig. 3C). The NfrA-like and Hypothetical Protein Type 2 have terminal aromatic residues and glycines at the -8/-10 and -7 position respectively. Other motifs show no similarity to prototypical β-signal motifs while many still show typical TMBB hydrophobicity/hydrophilicity alternation.

Finally, the protein types are distributed differently across phyla. We find TMBBs in 38 phyla and 88 classes. Prototypicals are found in all phyla and LptD-like proteins in most phyla. All TMBB types are found in at least some members of the Proteobacteria class and Firmicutes phylum and most are found in the Bacteroidetes phylum.

## Discussion

### Algorithm Benefits and Limitations

One of the difficulties of generalizing about the family of outer membrane β-barrels is the small number of solved crystal structures for proteins with low sequence similarity (6, 46, 47) With 1,894,206 sequences, our high-accuracy IsItABarrelDB database contributes a large set of sequences for analyzing TMBBs. The IsItABarrel algorithm can also be used to identify more TMBBs within genomes sequenced in the future.

In addition to IsItABarrelDB being at least two times larger than any previous TMBB database, the IsItABarrel algorithm selects between two and nine times fewer false positives (Table 1). Thus, the database is both larger and more reliable. However, the fact that IsItABarrel is more selective also results in a higher false negative rate. IsItABarrel incorrectly predicts 8% of TMBBs as non-TMBBs. The algorithms that have fewer false negatives have more false positives.

By relying on contact maps generated by homology, IsItABarrel has at its disposal a richer set of features than other algorithms. The other three predictors use independent discriminatory features such as hydrophobicity alternation in strand residues, average length of strands, detection of β-signal motif, differences in amino acid composition across the outer membrane and topology modeling with hidden Markov models. We speculate that the contact maps we use combine all of these data into a richer set of features that allows IsItABarrel to improve on prediction accuracy over previous methods. Thus, although the IsItABarrel 8% false negative rate shows there is room for future improvement, because the low false negative rate and high false positive rate are correlated for other algorithms and because the features used in other algorithms have implicit representation in IsItABarrel, it is unlikely that implementing features from other algorithms would help our algorithm reduce false negatives.

The two sources of error for our algorithm are the algorithm itself which has a balanced accuracy of 95.88% and the labeling assignments by sequence similarity > 40% which has accuracy of approximately 98% (Fig. 1F).

Though the contact maps from RaptorX are a key element of our algorithm, the main drivers of prediction accuracy are our features that describe strand-strand interactions and closing contacts. Therefore other contact map generation methods could have been employed, such as those from GREMLIN (35) or AlphaFold (36).

To assist with appropriate assessment of the impact of this inaccuracy in downstream tasks we provide annotation that identifies sequences that were predicted directly using IsItABarrel from those that had their predictions assigned by similarity (available online). Future analysis of the cases that receive different prediction labels when predicted by IsItABarrel and when being assigned by sequence similarity may reveal avenues for further improvement.

### TMBB by Phylogenetic Class

Using reference genomes, we find that previous reports of TMBB content being 2 – 3% of diderm bacterial genomes (48) are accurate for the most frequently studied of γ-proteobacteria genomes. However, over a larger range of organisms there is higher variance, from almost 7% to less than 1% with an overall mean of 1.54% and two modes at 0.4% and 1.4%. The category with the largest median genomic percentage of TMBBs is Bacteroidetes at a median of 3.34% per genome. It is possible that the greater number of TMBBs is due to Bacteroidetes localization in the gut where they need to utilize a variety of nutrients. For example Bacteroidetes use several polysaccharide utilization loci encoding outer membrane proteins that allow these organisms to adapt quickly to differences in the set of nutrient available. (49)

One notable feature is that some of the classes with the fewest numbers of TMBBs have the widest variety of TMBBs. Firmicutes, the second largest phylum found in the human microbiome after Bacteroidetes, has only a small number of organisms with TMBBs. This is likely due to the organisms in this phylum being low-GC Gram-positives with only some organisms being diderms with outer membranes. However, though Firmicutes tend to have few TMBBs as a percentage of their genome, they have the greatest variety of TMBBs along with Proteobacteria with both containing all ten types of TMBB. Bacteroidetes is a close third among the phyla with all but the NfrA-like proteins (Fig. 3D). Similarly, we find that several organisms in the Actinomycetia class have a small number of TMBBs, yet there is a large variety of TMBB types, with eight of the ten types of TMBBs in the phylum Actinobacteria.

### Domains of Unknown Function

We also analyzed the presence and distribution of DUFs in predicted TMBB sequences. In our reference genome analysis, we found 2,283 proteins that included domains labeled as DUF, which represents 5.21% of the set of new TMBB proteins found in these reference organisms. A total of 52 different folds categorized as DUFs are found among those 2,283 proteins (Table S4). For the whole IsItABarrelDB analysis we found 69,722 predicted TMBBs with DUFs (this represents 3.68% of our final dataset that contains a total of 1,894,206 sequences – Table S5). The distribution of sequences for each of the 280 DUFs follows a power law (Table S6), where only a few groups are large while most other groups have a much smaller number of sequences. In particular, we note that the two largest groups contain sequences that are mostly in the Prototypical type. The third group, DUF1302, formed a type of its own in our analysis (Fig. 3). In the reference genome analysis, we see that some of the domains we find have similarity to other barrels (DUF481, DUF3187) while others are less similar to whole barrels and may be a globular domain attached to a barrel protein and exposed on the cellular surface such as domains for cell adhesion (DUF1566).

### TMBB Sequence Length

The distribution of sequence lengths in the IsItABarrelDB is different across the ten protein types we identified (Fig. 3A). The largest type of Prototypical proteins is the most varied and shows a trimodal distribution where the peaks roughly indicate the possibility of doubling events from which outer membrane β-barrels may have evolved and increased in size by gene duplication (8, 12 and 16-stranded barrels).(3, 4) However, the Fim/Usher type is bimodal without showing doubling behavior. The smaller peak (242 residues) is likely indicative of an artifact due to the presence of partial sequences in our database. It is unlikely to be full barrels, since the proteins in this type have functions associated with protein translocation and we expected these barrels would have a large size. The type NfrA-like is also bimodal showing a possible doubling event. The types with the longest proteins, Fim/Usher, LptD-like and NfrA-like, have small “shoulders” indicating the possibility that these protein types have undergone barrel growth by a different mechanism, possibly hairpin addition, instead of a full doubling event.

### β-signal Motif

The vast majority of phylogenetic classes have the motif H_y_GH_y_GY+F, and there are variations on this theme that are organism dependent (Fig. 2C). Actinomycetia was unique among organism classes for having a LamB-like signal (W at -1, G at -7) dominate. (42) Though LamB itself is sufficiently related to prototypical barrels that they were typed together for this analysis, we see a LamB like β-signal for NfrA-like proteins and Tsx-like proteins, two types of proteins found in Actinomycetia (Fig. 3).

When looking at the β-signal by protein type we find that the class analysis was dominated by prototypical barrels. Moreover, though we analyzed the β-signal of the Fim/Usher proteins we did not find that it conformed to previous reports. (42) However, we do see the previously reported increase in histidine at the -3 position of β-proteobacteria (50)

Overall, there is little consensus among the β-signals when they are broken out into protein type. Prototypical and hypothetical type 2 maintain an aromatic at -1, G at -6. However, patterns among the others are less clear. Although we observe some positional conservation, hydrophobicity alternation, and the presence of an aromatic residue towards the end, a consistent motif is elusive. To attempt an evaluation of the consistency of the motif, we also analyzed the location where they were found (Figs. S3 and S4). While most motifs for our reference genome analysis were found at the end of the sequence, for the ten protein types the location is usually offset from the C-terminal end of the sequence and shows slightly more spread.

Our computational approach to detect consensus motifs may be influenced by sample size and sequence redundancy, which may have impacted detection of the true motif in some of the phyla with more variation. More studies are needed to understand the role of these motifs. However, because we see hydrophobicity alternation in all motifs, these motifs are likely localized within the barrel inserted in the membrane.

It is possible that the observed variations in the C-terminal motif, both in our phylogenetic class analysis and in our protein-type analysis, reflect different pathways of membrane insertion. These different types of proteins may use different C-terminal β-signals to interact with the Barrel Assembly Machinery (BAM), or some of these protein types may use different pathways for insertion. The β-signal also has a role in activating proteolysis machinery when unfolded barrels accumulate in the periplasm (51), and these differences in motifs may also be related to this additional function.

## Conclusion

We have developed an algorithm for TMBB prediction and have created a new database of curated TMBB sequences that is 40% more accurate and at least two times larger than those created with previous algorithms. TMBB content of bacterial genomes varies with some having more than 6% of the genome devoted to encoding TMBBs and some with less than 1%. For the most part we find that the β-signal varies in small ways by organism, and that the β-signal consensus is L/V-G-L/V-G-Y-R/K-F. A study of ten types of TMBB sequences indicates that there are many types of TMBBs without this consensus signal. We offer our TMBB identifying algorithm and database as a useful tool for researchers working with bacterial TMBBs who need a curated set of these proteins at https://isitabarrel.ku.edu andhttps://github.com/SluskyLab/isitabarrel. Awareness of the types and quantities of outer membrane proteins in different bacterial classes will be a foundation for better understanding of bacterial phylogeny/evolution and clearer understanding of outer membrane protein function.

## Materials and Methods

### IsItABarrel Algorithm

To classify sequences as TMBBs or non-TMBBs we combined RaptorX-Contact co-evolutionary contact map generation (52) with the development of a rule-based algorithm for feature extraction and classification. Map creation is more fully described in Supplemental Information, *Contact Map Generation*).

Our IsItABarrel algorithm was based on the following principles: Because the β-strands in TMBBs are ordered contiguously (strand 1 is next to strand 2, strand 2 is next to strand 3, etc.) TMBBs have consistent and distinctive contact map features. These features include map lines that indicate strand-strand interactions, and a line in the corner indicates the closing contacts between the first strand and the last strand. Non-TMBB folds rarely have both these features. The strand-strand contact feature is extracted by scanning the contact map along the main diagonal and counting how many strand-strand contact pairs are above a length threshold (further described in Supplemental Information, *Knowledge-Based Features and IsItABarrel Algorithm*). The closing contacts feature is extracted by scanning the corner regions of the map and ensuring the detected closing contacts align with previously found first and last strand. The two features are combined to classify the map as a TMBB or not-TMBB (Supplemental Information, *Knowledge-Based Features and the IsItABarrel Algorithm*). We observed that contact maps of sequences sharing more than 40% similarity receive the same IsItABarrel prediction 98% of the time. This made it possible to use IsItABarrel to predict large datasets in three steps: (1) clustering the set, (2) making a prediction for the smaller number of cluster representatives and (3) broadcasting this prediction to all other members of the cluster (further described in Supplemental information).

### Datasets

To train and validate the IsItABarrel algorithm we created three datasets of proteins with solved structures in the PDB. We call these datasets TMBB121 (Dataset S1), NONTMBB1K (Dataset S2) and SS1929 (Dataset S3). The TMBB121 dataset includes TMBB examples (TMBBs), while the sets NONTMBB1K and SS1929 include non-TMBB examples. We use only one set of TMBB examples since there is a limited number of non-redundant TMBBs with solved structures. The list of TMBB proteins were taken from a previously published dataset of outer membrane proteins (4) and the non-TMBB examples were randomly sampled from PISCES (53) and PDBSelect. (54) The negative examples were manually verified to ensure they did not include TMBBs. Since both of the datasets NONTMBB1K and SS1929 are much larger than the set of TMBB examples, we work with unbalanced data throughout our training, validation and testing. Dataset SS1929 is a dataset of proteins with signal sequence (as predicted by SignalP 5.0. (55)) (Supplemental Information, *Datasets)*. A summary of the datasets and how they were used is given in Table S1. The parameters of the IsItABarrel algorithm were optimized by grid search using balanced accuracy as a scoring metric on our validation set and details are provided in section *Lowering the False Negative Rate of IsItABarrel* in Supplemental Information and in Fig. S1).

To perform large-scale comparisons between different TMBB predictors (Fig. S2) we created two non-redundant datasets of protein sequences. The TMBB29183 (Dataset S4) contains the top 29,183 putative TMBB sequences from the TMBB-DB. (31) The OMPDB-NR (Dataset S5) contains 651,874 non-redundant representative sequences extracted from the OMPdb. Finally, using our IsItABarrel algorithm, we predicted TMBBs in the large non-redundant set of protein sequences from NCBI. This search resulted in the creation of our final IsItABarrelDB dataset with 1,894,206 TMBB sequences which is available for search and download at https://isitabarrel.ku.edu. Further details of how the database was created and features of our webapp are provided in section *Creation of the IsItABarrelDB* of the Supplemental Information.

To understand how TMBBs are distributed across taxonomic lineages, the proteomes of 2,959 prokaryotic organisms were analyzed with IsItABarrel. The sequences predicted to be TMBBs were counted by class/phylum (16 categories), producing updated estimates of TMBB content in prokaryotic organisms. We also used these sequences to observe differences in the distribution of lengths and C-terminal β-signal motif across the 16 taxonomic categories (Supplemental Information, *Distribution of TMBBs in Representative Organisms*).

To detect new types of TMBBs we performed a thorough sequence search of our final IsItABarrelDB dataset. Sequences of TMBBs with solved structures from all known TMBB types were used as seed queries against the IsItABarrelDB. Sequences that were not related to any of the known classes were clustered at 25%, and the largest clusters (designated ‘types’) were selected for analysis.

To understand how sequence length and β-signal motif are distributed across these new types of TMBBs we computed kernel density estimates (KDE) for the distribution of lengths of sequences and used MEME to compute motifs on the C-terminal quarter of the sequence Details of this analysis are given in Supplemental Information, A*nalysis of Sequences in IsItABarrelDB*. Details of the availability of our data and code are given in the Supplemental Information, *Code availability* and *Programming*.

## Acknowledgments

We thank Rachel Kolodny for important discussions about the algorithm, Harris Bernstein for helpful feedback on the manuscript, and Robert Unckless from the University of Kansas for help with our analysis of distribution of transmembrane β-barrels in prokaryotes. Financial support for this work came from National Institute of Health award DP2 GM128201 and P20GM103418, National Science Foundation (NSF) award 2226804, an American Scandinavian Foundation fellowship, and Kansas University (KU) Startup funding to Joanna Slusky. Daniel Montezano acknowledges support from the National Institute of Health award P20GM103418.

