## Supporting Information for "General Features of Transmembrane Beta Barrels From a Large Database"

#### **Supporting Information for**
#### General Features of Transmembrane Beta Barrels From a Large Database

Daniel Montezano, Rebecca Bernstein, Matthew M. Copeland, Joanna S. G. Slusky*

*Joanna S. G. Slusky.


**This PDF file includes:**

Supporting text
Figures S1 to S4
Tables S1 to S7
Legends for Datasets S1 to S5
Legend for Software S1

**Other supporting materials for this manuscript include the following:**

Datasets S1 to S5
Software S1

#### Supporting Information Text

##### Detailed Methods.

*Contact Map Generation*

Contact maps predicted by RaptorX are given by a symmetric matrix of numbers in the range [0,1], such that the upper triangular part above the main diagonal is identical to the lower triangle below the diagonal. RaptorX-Contact computes contact predictions for all possible pairs of residues in a sequence. However, only a fraction of the points on the map represents strong and confident predictions. To reduce the impact of weak predictions on our algorithm and improve the classification robustness we threshold the contact map, retaining only a predefined number of predictions proportional to the length of the sequence. For a sequence of length $L$ residues, in our tests with a small set of proteins, we observed that preserving the top strongest $2L*log_2(L)$ contacts improved the robustness of the classification without incurring significant loss of information that would preclude correct classification.

*Datasets*

Six different datasets of protein sequences were used in this work (Table S1). Due to the small number of non-redundant TMBB with solved structures available in the Protein Data Bank (PDB), there is only one dataset of known TMBB proteins with solved structures that is used throughout. The NONTMBB1K (Dataset S2) and SS1929 (Dataset S3) datasets include only non-TMBB sequences, all with solved structures in PDB. The sequence datasets TMBB29183 (Dataset S4) and OMPDB-NR (Dataset S4) were derived from the original databases (respectively the TMBB-DB and the OMPdb) and are non-redundant datasets used to compare different predictors (Fig. S2). The result of our search and evaluation is IsItABarrelDB, the largest set of predicted TMBBs.

We created three datasets for training IsItABarrel—one of TMBBs (TMBB121 – Dataset S1), one dataset of non-TMBB (NONTMBB1K –Dataset S2) with sequences that had solved structures in the PDB, and structurally resolved non-TMBBs that had a signal peptide (SS1929 –Dataset S3). For the TMBB set, sequences were obtained from a previously published (1) dataset of outer membrane proteins that was restricted to less than 60% similarity with CD-HIT (2). This set was smaller than all other sets because of the limited number of structurally resolved non-homologous TMBBs available in the protein data bank (PDB).

For the NONTMBB1K set, one thousand non-TMBB sequences were randomly selected from PDBSELECT100 (Nov. 18, 2017) (3). We removed all TMBBs by searching descriptions for any of the words "*outer, membrane, barrel, transport, channel, porin*" and manually removing from the set any potential TMBBs. A balanced dataset containing 100 TMBB sequences and a subset of 100 non-TMBB sequences from NONTMBB1K was used to develop our IsItABarrel algorithm.

We desired to test IsItABarrel on proteins containing a signal peptide that are not TMBBs since we planned to cull datasets first by removing proteins without a signal sequence. We developed the SS1929 dataset by randomly selecting 1,929 non-TMBB sequences from the non-redundant PISCES set cullpdb_pc20_res3.0_R1 (4). Selection continued until 1929 were found with the following four required characteristics: 1) a signal peptide by SignalP 5.0 (5), 2) overall sequence length between 120 and 1040, 3) match with a sequence in Uniprot, 4) no description of a possible barrel (as described above for the PDBSelect set).

All dataset sequences are available for download from https://github.com/SluskyLab/isitabarrel. Due to the large size of the set of contact map files (>3Tb) we are not able make them available in the same GitHub repository, but they are all available upon request. As of Jan. 2021, the original TMBB-DB and OMPdb datasets may be obtained from the webpages of the original authors at http://beta-barrel.tulane.edu/downloads.html and http://www.ompdb.org respectively.

*Knowledge-Based Features and the IsItABarrel Algorithm*

We used a grid search over a validation set (described in the next section) to identify ideal numerical parameters and thresholds (Fig. S1). Our IsItABarrel predictor extracts features from the generated contact maps that quantify the presence of a closing contact between the first and last strand and anti-parallel β-strand contacts. These two features are designated *strand-strand contact* and *closing contact*. These features are common in contact maps of TMBBs but not in contact map of non-TMBBs (Figs. 1A and 1B). Our *strand-strand contact* rule detects the number of lines perpendicular to the main diagonal as a proxy for the number of pairs of antiparallel β-strands interacting (Fig. 1C).

To find the contacts between two neighboring strands we consider 52x52 boxes going through the diagonal of the contact map and calculate the sum of the elements lying on the box diagonal that is perpendicular to the main map diagonal. The value of 52 for the box size (N Box, Fig. S1) was selected using a grid search over all parameters of IsItABarrel. To capture perpendicular lines that are near the corner of the map, we extend the map with zeros on all four sides before scanning the diagonal. We can graph the sums of the perpendicular lines by the residue number at the center of their scanning boxes (Fig. 1C). For any position where the sum is above a threshold of 18 (Z threshold, Fig. S1, with units of distance between points in the contact map), the area to the bottom left of the main diagonal is searched to locate the position of the perpendicular line that is closest to the main diagonal (perpendicular lines are often disconnected from the main diagonal). Then, the bottom left of the starting position is searched, and the number of residues between the starting position and the next strand–strand interaction position is summed. This sum is the length of the strand-strand contact. Additionally, to accurately compare β-hairpin lengths (two neighboring β-strands with a loop region connecting them is commonly called a "β-hairpin"), we calculate the length of the longest part of each β-hairpin by calculating the lengths of several lines along the hairpin and choosing the maximum value. Contact lengths between 16 and 25 residues (H4 low threshold and H4 high threshold, not shown) are defined as strand-strand contacts.

Our closing contact identification algorithm works in two steps. First, it detects all β-hairpins as previously described. Second, it iterates over the detected strands and then uses their location to search for the slanting line of closing contacts aligned to a pair of the first and last strands (Fig. 1D).

Once a potential closing contact is located, the position is saved and a 10x10 box (box size, Fig. S1) is created at the right upper corner of the map. If there is a full diagonal line from the left bottom corner to the right upper corner of the 10x10 box, the algorithm detects a closing contact. If not, a larger outline box with height 69 residues and width 7 residues (scan region, Fig. S1) is defined instead of the original 10x10 box. If a full diagonal line is found when scanning from the left bottom corner to the right upper corner in any 10x10 box inside this larger outline, it is defined as a closing contact.

The strand-strand contact prediction and closing contact prediction are then combined (Fig. 1E). If a closing contact is predicted, then the map is labeled a TMBB. In the absence of a detectable closing contact, if at least four strand-strand contacts are identified then the map is also labeled a TMBB. Otherwise, the map is defined as a non-TMBB.

*Lowering the False Negative Rate of IsItABarrel*

The IsItABarrel algorithm depends on various parameters and after developing the general method we optimized parameters to improve its accuracy (Fig. S1). To create a training set, we combined the full TMBB121 set with the full NONTMBB1K set into an unbalanced dataset containing 1,121 sequences and optimized the parameters of the algorithm by grid search. To lower the IsItABarrel false negative rate, during parameter refinement 21 TMBB sequences were selected based on visual inspection of contact maps that had TMBB signatures but were not being classified by IsItABarrel. Those 21 sequences were selected from 200 sequences that were identified as β-barrels by PRED-TMBB2 (6), BOMP (7), and the Freeman-Wimley (8) predictors, though not by initial parametrization of IsItABarrel.

*Creation of the IsItABarrelDB*

To create IsItABarrelDB, cluster representatives were obtained by clustering the initial set down to 30% similarity using MMSeqs2 (9) and predicting signal sequences with SignalP5.0. Contact maps for the set of 931,269 cluster representatives were generated with RaptorX-

Contact and predicted with IsItABarrel. Predictions from the cluster representatives were assigned to all other cluster members according to sequence similarity. Pairwise sequence similarity was determined using the stretcher algorithm from the EMBOSS suite (10). In this set, only sequences with length in the range of 120–1040 residues from bacterial organisms were included. The IsItABarrelDB dataset is the result of running IsItABarrel on the putative bacterial membrane proteins available in the large non-redundant protein set from NCBI (accessed March 2020).

The set of all sequences predicted with our IsItABarrel algorithm is available for search and download online at https://isitabarrel.ku.edu. In this database the user can select a subset of sequences from our resulting TMBB dataset by filtering the records using various available criteria, including the original Freeman-Wimley β-barrel score, the new IsItABarrel prediction and the SignalP 5.0 prediction. The database search can be refined further by selecting between the three different types of prokaryotic signal peptide pathways that SignalP 5.0 is able to distinguish: SP (the general secretory pathway with signal peptidase I), TAT (the twin-arginine translocation pathway with signal peptidase I, and LIPO (the translocation pathway for bacterial lipoproteins with cleavage by signal peptidase II). The user can also search by organism using partial matching of boolean expression (e.g. *coli* AND *pestis*) and by sequence length. The database also includes other information such as the protein type the entry belongs to and whether the IsItABarrel prediction has been computed from the map or assigned by similarity from the cluster representative. The sequence representative used for assignment is also provided. Currently, the filtered results are generated, and then an email is sent to the user with a link to download the results of the search as a compressed tar archive file. The archive contains two individual data files and the license file. The first data file is a records file containing the returned records in TAB-separated format. The second data file is a FASTA file containing the amino acid sequences of the corresponding records with the non-redundant protein accession number as the FASTA header to use as a key between the two tables.

*Distribution of TMBBs in Representative Organisms*

To compute the distribution of TMBB populations in prokaryotic organisms, we searched bacterial genome assemblies of 2,944 representative organisms and 15 reference organisms for TMBB homologs. Genomes were retrieved from the NCBI Assembly database (April 2021) using filters "Latest", "RefSeq", "Complete Genome" and "Exclude anomalous". Homologs were found between the 95% sequence-similarity-filtered OMPDB-NR set and the RefSeq proteomes using blastp from BLAST version 2.9.0+ (11), E-value of 1.0, query coverage of 95%, and all other parameters at their default values. We removed any redundant hits and filtered the results of each search by retaining only hits with 90% or more positive matching residues. Different TMBBs matching the same protein sequence were counted only once.

To compute the distribution of TMBBs per organism we grouped the 2,959 organisms by taxonomic class with the exception of Bacteroidetes and Firmicutes organisms which were grouped by phylum.

*Analysis of Sequences in IsItABarrelDB*

To identify types of proteins from the whole IsItABarrelDB we used sequences from a previously published dataset of structurally characterized TMBBs separated by homology in different TMBB categories (1). We used the sequences of the structurally solved proteins as search queries to find homologs in the IsItABarrelDB using MMSeqs2 with an E-value of 1E-12. The matches to each of the categories were used as queries against the remaining part of the IsItABarrelDB. Only significant matches were considered that satisfied two criteria: (1) minimum alignment length of 250 residues and (2) coverage of at least 75% for either query or target whichever was shorter. This stringent cumulative search was performed for six iterations allowing us to cluster a total of 1,640,974 sequences. During this iterative search we also considered overlap, merging groups of search targets when the same sequence matched in different categories. From the clustered sequences we derived five types of TMBB sequences: *Prototypicals*, *Fim/Usher*, *LptD-like*, *LpxR-like* and *Tsx-like*. We collected the remaining 253,232 sequences in the unrelated remainder. We clustered the unrelated remainder at 25% using MMSeqs2 and used the five clusters with more than 3,000 members for downstream analysis.

We named these protein types according to information retrieved from their FASTA headers: *DUF1302*, *NfrA-like*, *PorT-like*, *Hypothetical Protein Type 1* and *Hypothetical Protein Type 2*. We note that by using only the largest clusters we still left out 232,840 sequences (unclassified) from our final analysis. For each of the final ten types (five related to structurally resolved barrels, five unrelated to structurally resolved barrels) we computed the distribution of lengths and their distribution across bacterial phyla. For the MEME β-signal motif search we used the last quarter of each sequence (closest to the C-terminus). Location of the motifs found are given in Figs. S3 and S4.

*Code availability*

Code for RaptorX is freely available for local installation from the original authors and can be downloaded at https://github.com/j3xugit/RaptorX-Contact. All parameters were left at their defaults, but differences in contact map generation may arise due to the differences in the underlying databases required by the software. Predictor PRED-TMBB2 is available as a webapp only (http://www.compgen.org/tools/PRED-TMBB2), BOMP is available only as a webapp (http://services.cbu.uib.no/tools/bomp), and the Freeman-Wimley Beta-Barrel predictor is available only as a Windows executable (http://beta-barrel.tulane.edu/downloads.html).

*Programming*

All steps of dataset creation and analysis were implemented with custom BASH shell and Python scripts specific for use with the KU High-Performance cluster, using the parameters given in the supplemental material. Contact map estimation using RaptorX was also performed on the high-performance computer cluster at the University of Kansas.

*Parameters*

The parameters used for ***(A)*** *finding queries used for NCBI search of prokaryotic reference genomes,* ***(B)*** *performing blast search against known TMBBs;* ***(C)*** *performing MEME motif search (MEME version 5.3.3)* ***(D)*** *performing MMSeqs2 clustering,* ***(E)*** *performing CD-HIT clustering,* ***(F)*** *generation of kernel density estimates, and* ***(G)*** *computation of pairwise sequence similarity:*

(A) prokaryota[orgn] AND “representative genome”[refseq category]
prokaryota[orgn] AND “reference genome"[refseq category]

(B) **makeblastdb** -in <PROTEOME.faa> -dbtype prot -out <PROTEOME_DB>
**blastp** -db <PROTEOME_DB> -query <INPUT_KNWON_TMBBS.faa> -task blastp -num_threads 16 -evalue 1.0 -qcov_hsp_perc 95 -outfmt “6 qaccver qlen saccver slen evalue length pident ppos”

(C) **meme** <INPUT.faa> -nostatus -oc <OUTPUT_FOLDER> -protein -mod oops -nmotifs 1 -w 8

(D) # format database and cascade clustering at 30%
**mmseqs** createdb <INPUT.faa> DB
**mmseqs** cluster DB DB_clu30 tmpfolder/ --min-seq-id 0.3
### create fasta file with representatives
**mmseqs** createsubdb DB_clu30 DB DB_clu30_rep
**mmseqs** convert2fasta DB_clu30_rep <REPRESENTATIVES.faa>
### create mapping between representatives and members
**mmseqs** createtsv DB DB DB_clu30 <MAPPING.tsv>

(E) cd-hit -i <INPUT.fasta> -o <OUTPUT> -c 0.6
All other CD-HIT parameters at their defaults.

(F) kernel density estimates of sequence lengths were computed with the R function **density()**, with 512 points each or with the function smooth in Gnuplot 5.4.2.

(G) **stretcher** -asequence <SEQA.fasta> -bsequence <SEQB.fasta> -sprotein1 \
-sprotein2 -sformat1 fasta -sformat2 fasta -gapopen 12 -gapextend 2 \
-datafile EBLOSUM62 -stdout -auto
and extracting from the resulting file the percentage in the line “#Similarity”.

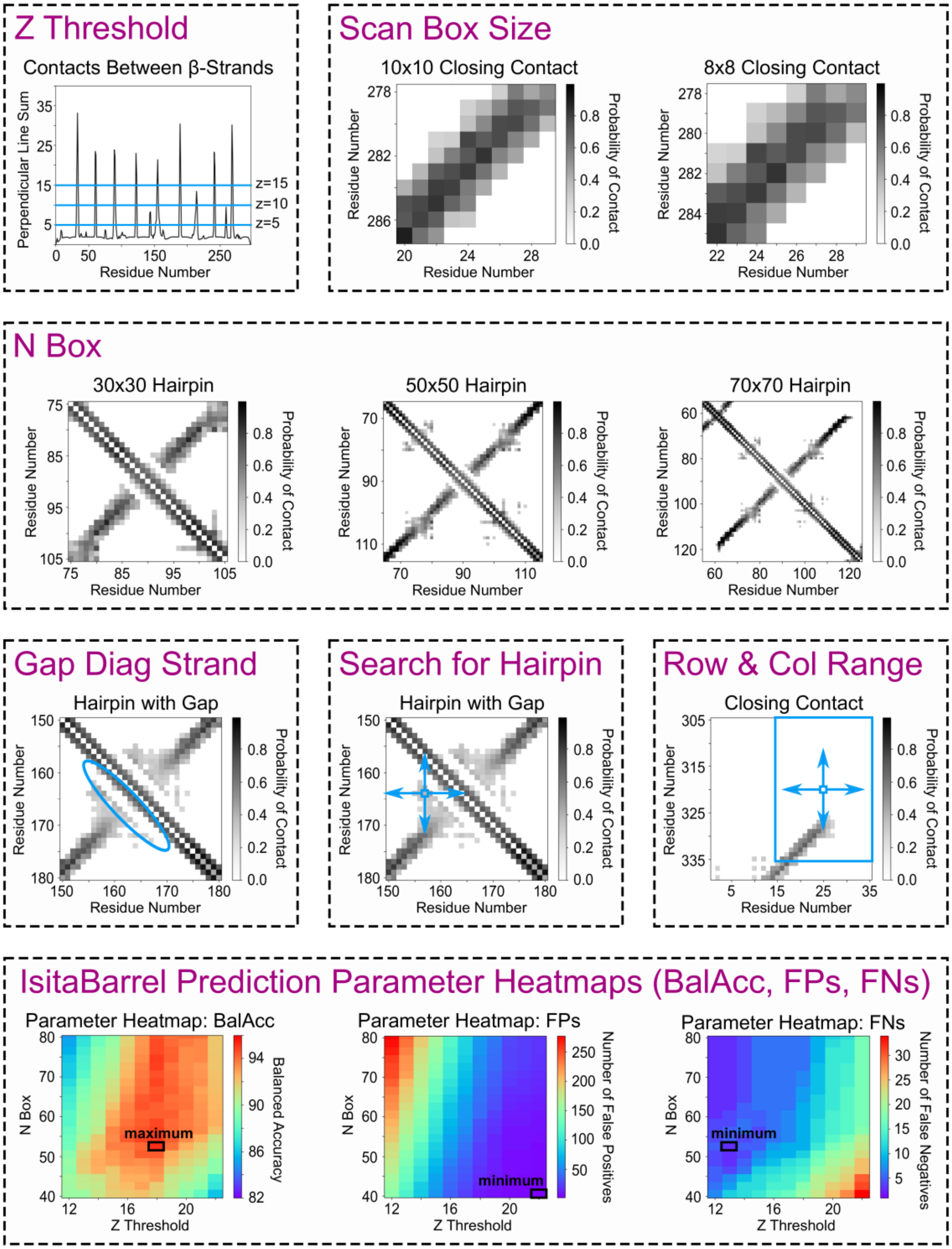


**Figure S1. The Parameters of the IsItABarrel Algorithm.** Top three rows show some of the different parameters for the IsItABarrel algorithm: Z Threshold (threshold in which a peak is counted as a hairpin), Scan Box Size (size of box when scanning for closing contact), N Box (size of box when scanning for hairpins), Gap Diag Strand (maximum number of residues to go out past middle diagonal when searching for start of hairpin to account for possible gap between hairpin and middle diagonal), Search for Hairpin (maximum number of residues to go out from presumed start of hairpin in four directions to find a new start of hairpin that starts at the point where hairpin has the largest length), Row Range (width of box created around residue that was pinpointed for closing contact), and Column Range (length of box created around residue that was pinpointed for closing contact). Bottom row shows three parameter heatmaps created after running 451 simulations in which Z Threshold and N Box are changed simultaneously. For each simulation, the NONTMBB1K with TMBB121 dataset is used to test the balanced accuracy, number of false positives, and number of false negatives of the IsItABarrel algorithm for different combinations of Z Threshold and N Box. These three parameter heatmaps show the results of the simulations: the first heatmap shows the balanced accuracy of the IsItABarrel algorithm for different combinations of Z Threshold and N Box; the second heat map shows the number of false positives for different combinations of Z Threshold and N Box; the third heatmap shows the number of false positives for different combinations of Z Threshold and N Box.

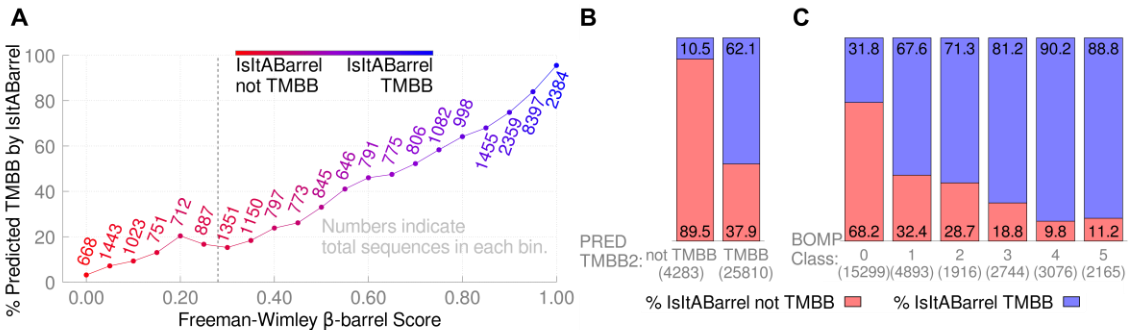


**Figure S2.** Comparison of IsItABarrel with Three Other TMBB predictors. **(A)** For the Freeman-Wimley algorithm a β-barrel score is computed, and the sequence is classified as a TMBB for a score above 0.28. β-barrel scores for a set of 29,183 sequences were grouped in bins of width 0.05. In each bin IsItABarrel detects a percentage of barrels, which decreases with lower Freeman-Wimley score, showing a correlation between the two algorithms. **(B)** Two-class prediction by PRED-TMBB2. There is significant overlap for non-TMBB predictions, while for TMBB predictions IsItABarrel finds 37.9% of BOMP predictions to be false positives. **(C)** The BOMP predictions for TMBBs also show significant overlap with IsItABarrel. For non-TMBBs (BOMP class 0), IsItABarrel identifies many false negatives (31.8% in class 0). We classified two non-redundant datasets OMPDB-NR and the TMBB29183 (Table S1) using each of the algorithms. Though it is not possible to derive conclusions about the absolute prediction accuracy of each algorithm since we do not have ground truth labels for the datasets, we find that all three methods correlate with IsItABarrel. For sets with ground truth labels (Table 1), the other algorithms all substantially over-predict barrels. We find that while the other two classifiers over-predict TMBBs, BOMP under-predicts barrels. BOMP classifies a putative TMBB sequence in one of six classes, of which classes four and five indicate a strong TMBB prediction and class zero indicates a negative prediction (not a TMBB). There is strong correlation in positive BOMP classes four and five. However, 31.8% of non-TMBB predictions (class zero) by BOMP are considered to be false negatives by IsItABarrel.

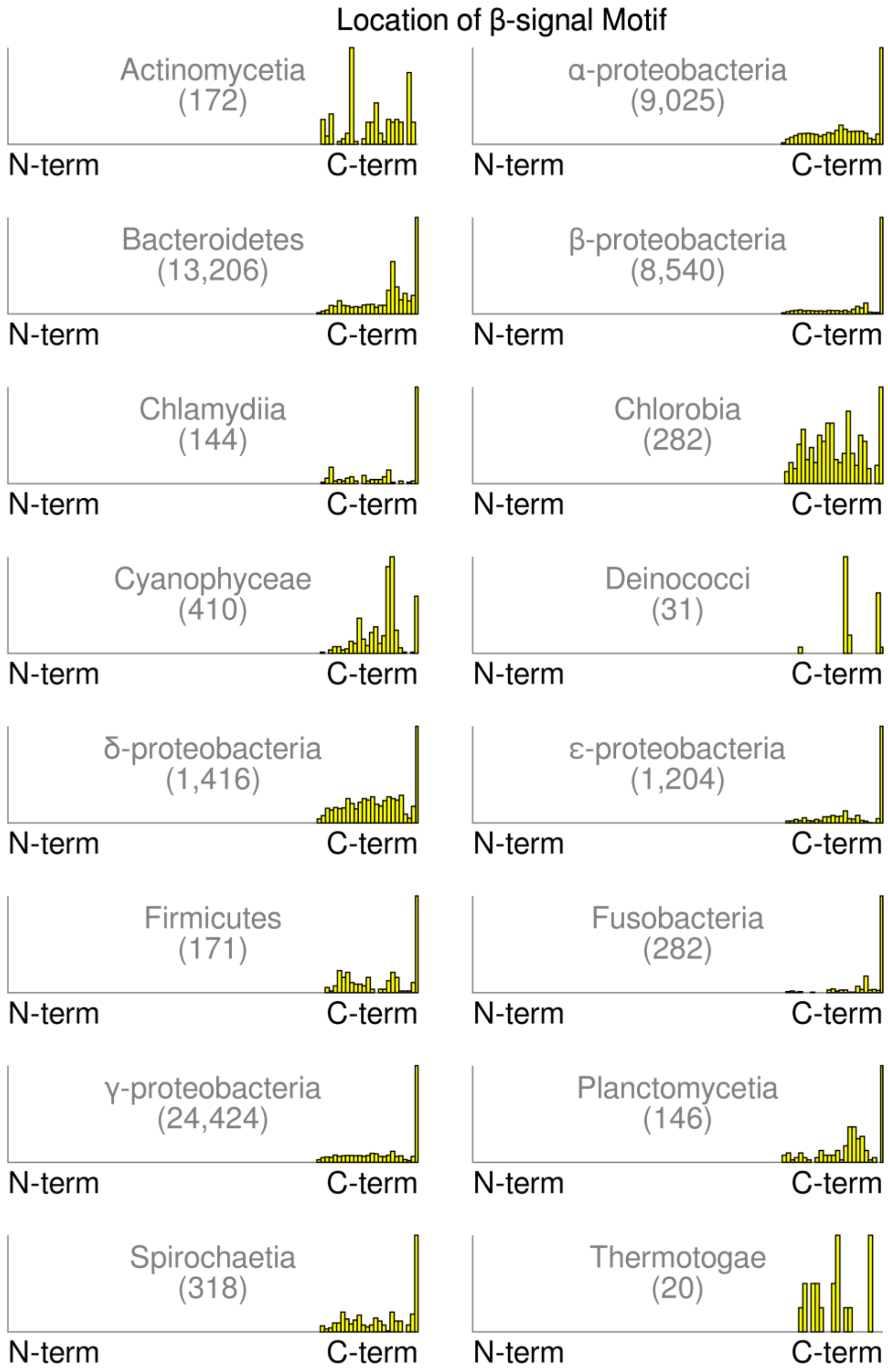


**Figure S3. β-signal Motif Locations for 16 Taxonomic Classes.** Histograms of the location of sequence motifs searched in the quarter closest to the C-terminal of each sequence using MEME. The x-axis shows relative distance along the full-length sequence.

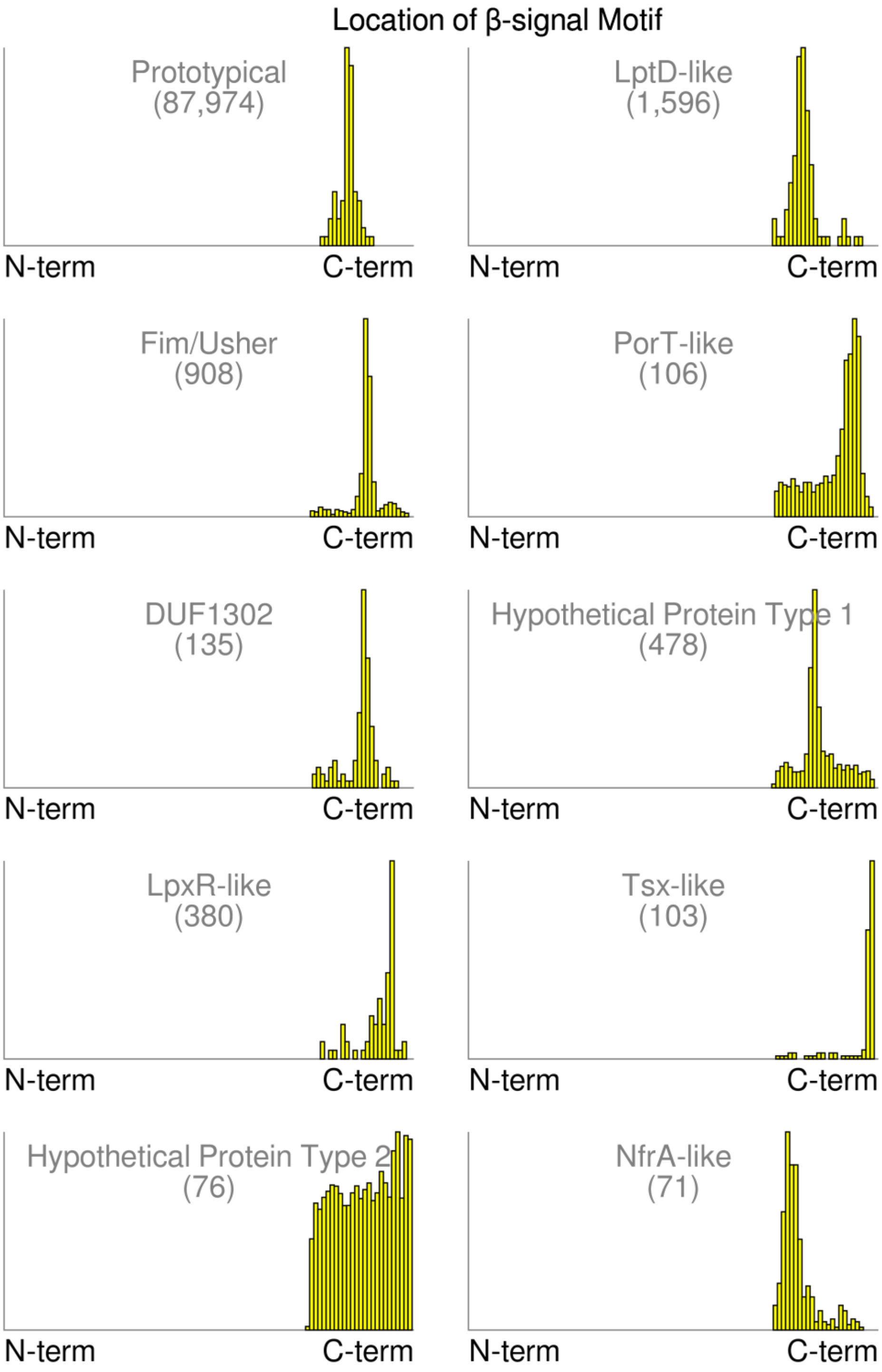


**Figure S4. β-signal Motif Locations for Ten Largest IsItABarrelDB Types.** Sequence motifs searched in the quarter closest to the C-terminal of each sequence using MEME. The x-axis shows relative distance along the full-length sequence.

**Table S1.** Datasets used to develop, test and apply IsItABarrel.

| Name | Description | Size | Usage |
|---|---|---|---|
| *TMBB121* | *Non-redundant TMBBs with solved structures.* | *121* | *Algorithm development, validation, comparison of predictors, distribution of lengths* |
| *NONTMBB1K* [a] | *Non homologous non-TMBBs with solved structures.* | *1,000* | *Parameter optimization* |
| *SS1929* [a] | *Non-homologous membrane non-TMBBs with solved structures.* | *1,929* | *Validation* |
| *TMBB29183* | *Non-redundant set of sequences from the TMBB-DB.* | *29,183* | *Comparison of predictors* |
| *OMPDB-NR* | *Non-redundant set of sequences from the OMPdb.* | *651,874* | *Comparison of predictors* |
| *IsItABarrelDB* | *Non-redundant set of bacterial sequences from NCBI NR database predicted TMBB.* | *1,894,206* | *Distribution of lengths* |

[a] In the text, whenever the sets NONTMBB1K and SS1929 were used, it is assumed that these are the negative examples, while the TMBB121 set was used in all cases as the set of positive examples.

**Table S2.** The 78 classes covered by the 2,959 representative and reference organisms used in the search for TMBBs.

| | | | | | |
|---|---|---|---|---|---|
| 1 | Acidimicrobiia | 27 | Deferribacteres | 53 | Mollicutes |
| 2 | Acidithiobacillia | 28 | Dehalococcoidia | 54 | Negativicutes |
| 3 | Acidobacteriia | 29 | Deinococci | 55 | Nitriliruptoria |
| 4 | Actinomycetia | 30 | δ-proteobacteria | 56 | Nitrososphaeria |
| 5 | α-proteobacteria | 31 | Dictyoglomia | 57 | Nitrospira |
| 6 | Anaerolineae | 32 | Elusimicrobia | 58 | Oligoflexia |
| 7 | Aquificae | 33 | Endomicrobia | 59 | Opitutae |
| 8 | Archaeoglobi | 34 | ε-proteobacteria | 60 | Phycisphaerae |
| 9 | Bacilli | 35 | Erysipelotrichia | 61 | Planctomycetia |
| 10 | Bacteroidia | 36 | Fimbriimonadia | 62 | Rubrobacteria |
| 11 | β-proteobacteria | 37 | Flavobacteriia | 63 | Saprospiria |
| 12 | Blastocatellia | 38 | Fusobacteriia | 64 | Sphingobacteriia |
| 13 | Caldilineae | 39 | γ-proteobacteria | 65 | Spirochaetia |
| 14 | Caldisericia | 40 | Gemmatimonadetes | 66 | Synergistia |
| 15 | Calditrichae | 41 | Gloeobacteria | 67 | Tepidiformia |
| 16 | Chitinophagia | 42 | Halobacteria | 68 | Thermococci |
| 17 | Chlamydiia | 43 | Hydrogenophilalia | 69 | Thermodesulfobacteria |
| 18 | Chlorobia | 44 | Ignavibacteria | 70 | Thermoleophilia |
| 19 | Chloroflexia | 45 | Kiritimatiellae | 71 | Thermomicrobia |
| 20 | Chrysiogenetes | 46 | Ktedonobacteria | 72 | Thermoplasmata |
| 21 | Clostridia | 47 | Limnochordia | 73 | Thermoprotei |
| 22 | Conexivisphaeria | 48 | Methanobacteria | 74 | Thermotogae |
| 23 | Coprothermobacteria | 49 | Methanococci | 75 | Tissierellia |
| 24 | Coriobacteriia | 50 | Methanomicrobia | 76 | Verrucomicrobiae |
| 25 | Cyanophyceae | 51 | Methanopyri | 77 | Vicinamibacteria |
| 26 | Cytophagia | 52 | Methylacidiphilae | 78 | Zetaproteobacteria |

**Table S3.** The 35 phyla covered by the 2,959 representative and reference organisms used in the search for TMBBs.

| | | | | | |
|---|---|---|---|---|---|
| 1 | Acidobacteria | 13 | Coprothermobacterota | 25 | Kiritimatiellaeota |
| 2 | Actinobacteria | 14 | Crenarchaeota | 26 | Nitrospirae |
| 3 | Aquificae | 15 | Cyanobacteria | 27 | Planctomycetes |
| 4 | Armatimonadetes | 16 | Deferribacteres | 28 | Proteobacteria |
| 5 | Bacteroidetes | 17 | Deinococcus-Thermus | 29 | Spirochaetes |
| 6 | Caldiserica | 18 | Dictyoglomi | 30 | Synergistetes |
| 7 | Calditrichaeota | 19 | Elusimicrobia | 31 | Tenericutes |
| 8 | Candidatus Thermoplasmatota | 20 | Euryarchaeota | 32 | Thaumarchaeota |
| 9 | Chlamydiae | 21 | Firmicutes | 33 | Thermodesulfobacteria |
| 10 | Chlorobi | 22 | Fusobacteria | 34 | Thermotogae |
| 11 | Chloroflexi | 23 | Gemmatimonadetes | 35 | Verrucomicrobia |
| 12 | Chrysiogenetes | 24 | Ignavibacteriae | - - - | - - - |

**Table S4.** The 52 domains of unknown function (DUF) found in predicted TMBBs of the 2,959 representative and reference organisms included in our reference genome analysis.

| | | | | | | | |
|---|---|---|---|---|---|---|---|
| **1** | DUF11 | **14** | DUF1302 | **27** | DUF2219 | **40** | DUF3999 |
| **2** | DUF370 | **15** | DUF1396 | **28** | DUF2442 | **41** | DUF4105 |
| **3** | DUF481 | **16** | DUF1488 | **29** | DUF2808 | **42** | DUF4139 |
| **4** | DUF756 | **17** | DUF1513 | **30** | DUF3108 | **43** | DUF4163 |
| **5** | DUF839 | **18** | DUF1540 | **31** | DUF3187 | **44** | DUF4331 |
| **6** | DUF916 | **19** | DUF1549 | **32** | DUF3298 | **45** | DUF4381 |
| **7** | DUF945 | **20** | DUF1553 | **33** | DUF3458 | **46** | DUF4403 |
| **8** | DUF1036 | **21** | DUF1566 | **34** | DUF3459 | **47** | DUF4440 |
| **9** | DUF1045 | **22** | DUF1573 | **35** | DUF3494 | **48** | DUF4595 |
| **10** | DUF1080 | **23** | DUF1579 | **36** | DUF3570 | **49** | DUF4625 |
| **11** | DUF1104 | **24** | DUF2135 | **37** | DUF3734 | **50** | DUF4982 |
| **12** | DUF1207 | **25** | DUF2147 | **38** | DUF3828 | **51** | DUF5050 |
| **13** | DUF1259 | **26** | DUF2191 | **39** | DUF3943 | **52** | DUF5103 |

**Table S5.** Predicted TMBBs with domains of unknown function (DUF) by organismal category in the 2,959 representative and reference organisms.

| Organismal category | How many proteins? | Which DUFs? |
|---|---|---|
| Actinomycetia | 159 | DUF1259, DUF1396, DUF2191, DUF3298, DUF3459, DUF4139, DUF4331, DUF4982, DUF756, DUF839, DUF916 |
| α-proteobacteria | 246 | DUF1036, DUF1045, DUF11, DUF1259, DUF1302, DUF1579, DUF3108, DUF3459, DUF4139, DUF4163, DUF4331, DUF4403, DUF4440, DUF481, DUF4982, DUF756, DUF839 |
| Bacteroidetes | 723 | DUF1080, DUF1573, DUF2135, DUF2147, DUF2219, DUF3458, DUF3459, DUF3494, DUF3570, DUF3828, DUF3943, DUF3999, DUF4139, DUF4163, DUF4331, DUF4403, DUF4440, DUF4595, DUF4625, DUF481, DUF4982, DUF5050, DUF5103, DUF756, DUF839 |
| β-proteobacteria | 105 | DUF1259, DUF1302, DUF1579, DUF2135, DUF3570, DUF4105, DUF4139, DUF4331, DUF4982, DUF756, DUF839, DUF945 |
| Chlamydiia | 1 | DUF1207 |
| Chlorobia | 0 | - - - - - - |
| Cyanophyceae | 50 | DUF1579, DUF2808, DUF3459, DUF4331, DUF839 |
| Deinococci | 46 | DUF11, DUF3459, DUF4403, DUF839 |
| δ-proteobacteria | 42 | DUF1259, DUF1302, DUF1549, DUF1573, DUF1579, DUF2219, DUF3187, DUF3570, DUF4105, DUF4139, DUF4163, DUF4331, DUF4381, DUF4403, DUF839 |
| ε-proteobacteria | 19 | DUF1104, DUF3570, DUF4105, DUF4139, DUF839 |
| Firmicutes | 100 | DUF11, DUF1259, DUF1540, DUF3298, DUF370, DUF3999, DUF4163, DUF4982, DUF839 |
| Fusobacteriia | 6 | DUF3298, DUF3999, DUF4163 |
| γ-proteobacteria | 352 | DUF1259, DUF1302, DUF1488, DUF1513, DUF1566, DUF1579, DUF2135, DUF2219, DUF2442, DUF3187, DUF3459, DUF3570, DUF3734, DUF3999, DUF4105, DUF4139, DUF4163, DUF4331, DUF4403, DUF4982, DUF756, DUF839, DUF945 |
| Planctomycetia | 338 | DUF11, DUF1549, DUF1553, DUF1573, DUF1579, DUF4139, DUF4331, DUF4982, DUF839 |
| Spirochaetia | 16 | DUF1566, DUF3187, DUF3298, DUF3943, DUF4105, DUF4139, DUF4163, DUF839 |
| Thermotogae | 5 | DUF4163 |

**Table S6.** The 280 domains of unknown function (DUF) found in predicted TMBBs in the IsItABarrelDB dataset by decreasing order of frequency. In parentheses is the count of the number of sequences annotated with each DUF.

| | Domain (count) | | Domain (count) | | Domain (count) | | Domain (count) |
|---|---|---|---|---|---|---|---|
| **1** | DUF1573 (10577) | **71** | DUF4082 (53) | **141** | DUF4989 (8) | **211** | DUF928 (1) |
| **2** | DUF4982 (9149) | **72** | DUF1396 (50) | **142** | DUF4480 (8) | **212** | DUF805 (1) |
| **3** | DUF1302 (8511) | **73** | DUF4412 (46) | **143** | DUF4424 (8) | **213** | DUF732 (1) |
| **4** | DUF3575 (7906) | **74** | DUF1735 (46) | **144** | DUF4419 (8) | **214** | DUF554 (1) |
| **5** | DUF560 (5872) | **75** | DUF4974 (45) | **145** | DUF4038 (7) | **215** | DUF5104 (1) |
| **6** | DUF4105 (4528) | **76** | DUF2012 (45) | **146** | DUF4623 (6) | **216** | DUF5024 (1) |
| **7** | DUF481 (4216) | **77** | DUF4906 (41) | **147** | DUF3471 (6) | **217** | DUF5010 (1) |
| **8** | DUF3373 (4125) | **78** | DUF4855 (41) | **148** | DUF1311 (6) | **218** | DUF5008 (1) |
| **9** | DUF5103 (3800) | **79** | DUF3472 (37) | **149** | DUF799 (5) | **219** | DUF4988 (1) |
| **10** | DUF3298 (3747) | **80** | DUF3769 (33) | **150** | DUF5006 (5) | **220** | DUF4961 (1) |
| **11** | DUF4163 (3388) | **81** | DUF3494 (32) | **151** | DUF4886 (5) | **221** | DUF4929 (1) |
| **12** | DUF3187 (3369) | **82** | DUF4430 (31) | **152** | DUF4838 (5) | **222** | DUF4785 (1) |
| **13** | DUF3078 (3185) | **83** | DUF2380 (29) | **153** | DUF4369 (5) | **223** | DUF4469 (1) |
| **14** | DUF3570 (3066) | **84** | DUF1597 (29) | **154** | DUF4360 (5) | **224** | DUF4366 (1) |
| **15** | DUF3108 (3036) | **85** | DUF3459 (28) | **155** | DUF3794 (5) | **225** | DUF4340 (1) |
| **16** | DUF2490 (2975) | **86** | DUF4384 (27) | **156** | DUF3179 (5) | **226** | DUF433 (1) |
| **17** | DUF916 (2588) | **87** | DUF2320 (26) | **157** | DUF1559 (5) | **227** | DUF4292 (1) |
| **18** | DUF5020 (2146) | **88** | DUF839 (23) | **158** | DUF882 (4) | **228** | DUF4271 (1) |
| **19** | DUF2219 (1412) | **89** | DUF4215 (23) | **159** | DUF5018 (4) | **229** | DUF427 (1) |
| **20** | DUF3138 (1352) | **90** | DUF4115 (23) | **160** | DUF4976 (4) | **230** | DUF4255 (1) |
| **21** | DUF3943 (1254) | **91** | DUF3280 (23) | **161** | DUF4652 (4) | **231** | DUF4200 (1) |
| **22** | DUF4139 (1033) | **92** | DUF5068 (22) | **162** | DUF4252 (4) | **232** | DUF4199 (1) |
| **23** | DUF11 (917) | **93** | DUF5019 (22) | **163** | DUF4249 (4) | **233** | DUF4168 (1) |
| **24** | DUF945 (897) | **94** | DUF4091 (22) | **164** | DUF377 (4) | **234** | DUF4101 (1) |
| **25** | DUF3324 (883) | **95** | DUF1800 (21) | **165** | DUF3659 (4) | **235** | DUF4073 (1) |
| **26** | DUF1566 (803) | **96** | DUF3238 (20) | **166** | DUF3573 (4) | **236** | DUF4012 (1) |
| **27** | DUF4403 (656) | **97** | DUF5115 (18) | **167** | DUF3442 (4) | **237** | DUF400 (1) |
| **28** | DUF5110 (600) | **98** | DUF1983 (18) | **168** | DUF3225 (4) | **238** | DUF3872 (1) |
| **29** | DUF1207 (570) | **99** | DUF1551 (18) | **169** | DUF303 (4) | **239** | DUF3859 (1) |
| **30** | DUF1036 (527) | **100** | DUF1410 (18) | **170** | DUF2141 (4) | **240** | DUF3826 (1) |
| **31** | DUF1553 (488) | **101** | DUF4381 (17) | **171** | DUF2125 (4) | **241** | DUF378 (1) |
| **32** | DUF4421 (475) | **102** | DUF4331 (17) | **172** | DUF1565 (4) | **242** | DUF362 (1) |
| **33** | DUF1579 (424) | **103** | DUF1554 (17) | **173** | DUF1416 (4) | **243** | DUF3607 (1) |
| **34** | DUF3316 (410) | **104** | DUF1202 (17) | **174** | DUF1194 (4) | **244** | DUF3568 (1) |
| **35** | DUF4968 (375) | **105** | DUF5121 (16) | **175** | DUF1058 (4) | **245** | DUF3500 (1) |
| **36** | DUF2860 (356) | **106** | DUF5107 (16) | **176** | DUF5126 (3) | **246** | DUF3440 (1) |
| **37** | DUF2808 (314) | **107** | DUF459 (16) | **177** | DUF4874 (3) | **247** | DUF3427 (1) |
| **38** | DUF4625 (300) | **108** | DUF4474 (16) | **178** | DUF4431 (3) | **248** | DUF3387 (1) |
| **39** | DUF58 (276) | **109** | DUF3604 (16) | **179** | DUF4394 (3) | **249** | DUF3383 (1) |
| **40** | DUF4979 (246) | **110** | DUF4347 (15) | **180** | DUF4179 (3) | **250** | DUF3374 (1) |
| **41** | DUF3308 (236) | **111** | DUF3808 (15) | **181** | DUF4129 (3) | **251** | DUF3352 (1) |
| **42** | DUF4165 (207) | **112** | DUF5060 (14) | **182** | DUF4124 (3) | **252** | DUF333 (1) |
| **43** | DUF3857 (207) | **113** | DUF5005 (14) | **183** | DUF4114 (3) | **253** | DUF3237 (1) |
| **44** | DUF2066 (203) | **114** | DUF4104 (14) | **184** | DUF3836 (3) | **254** | DUF3124 (1) |
| **45** | DUF4440 (187) | **115** | DUF4876 (13) | **185** | DUF2268 (3) | **255** | DUF3011 (1) |
| **46** | DUF1239 (168) | **116** | DUF3244 (13) | **186** | DUF1939 (3) | **256** | DUF2968 (1) |
| **47** | DUF748 (167) | **117** | DUF3068 (13) | **187** | DUF1593 (3) | **257** | DUF2911 (1) |
| **48** | DUF4382 (164) | **118** | DUF940 (12) | **188** | DUF1555 (3) | **258** | DUF2889 (1) |
| **49** | DUF3520 (154) | **119** | DUF756 (12) | **189** | DUF1192 (3) | **259** | DUF2817 (1) |
| **50** | DUF2807 (139) | **120** | DUF4981 (12) | **190** | DUF861 (2) | **260** | DUF2804 (1) |
| **51** | DUF4832 (135) | **121** | DUF4493 (12) | **191** | DUF5320 (2) | **261** | DUF2712 (1) |
| **52** | DUF3616 (126) | **122** | DUF4214 (12) | **192** | DUF461 (2) | **262** | DUF2531 (1) |
| **53** | DUF4198 (125) | **123** | DUF2006 (12) | **193** | DUF4434 (2) | **263** | DUF2436 (1) |
| **54** | DUF5117 (119) | **124** | DUF1906 (12) | **194** | DUF4344 (2) | **264** | DUF2394 (1) |
| **55** | DUF5011 (118) | **125** | DUF5016 (11) | **195** | DUF4232 (2) | **265** | DUF2334 (1) |

| | | | | | | | |
|---|---|---|---|---|---|---|---|
| **56** | DUF3999 (118) | **126** | DUF4990 (11) | **196** | DUF4175 (2) | **266** | DUF2300 (1) |
| **57** | DUF4397 (110) | **127** | DUF4962 (11) | **197** | DUF348 (2) | **267** | DUF2285 (1) |
| **58** | DUF1080 (110) | **128** | DUF3823 (11) | **198** | DUF3233 (2) | **268** | DUF2135 (1) |
| **59** | DUF3034 (108) | **129** | DUF2341 (11) | **199** | DUF323 (2) | **269** | DUF1812 (1) |
| **60** | DUF5686 (102) | **130** | DUF1083 (11) | **200** | DUF3224 (2) | **270** | DUF1794 (1) |
| **61** | DUF4892 (96) | **131** | DUF547 (10) | **201** | DUF2961 (2) | **271** | DUF1778 (1) |
| **62** | DUF1549 (83) | **132** | DUF5123 (10) | **202** | DUF285 (2) | **272** | DUF1775 (1) |
| **63** | DUF5050 (73) | **133** | DUF5074 (10) | **203** | DUF255 (2) | **273** | DUF1682 (1) |
| **64** | DUF5125 (69) | **134** | DUF4880 (10) | **204** | DUF2233 (2) | **274** | DUF1641 (1) |
| **65** | DUF1513 (69) | **135** | DUF4595 (9) | **205** | DUF1961 (2) | **275** | DUF1592 (1) |
| **66** | DUF3996 (67) | **136** | DUF4465 (9) | **206** | DUF1928 (2) | **276** | DUF1587 (1) |
| **67** | DUF5116 (60) | **137** | DUF4185 (9) | **207** | DUF187 (2) | **277** | DUF1486 (1) |
| **68** | DUF4959 (60) | **138** | DUF4140 (9) | **208** | DUF1768 (2) | **278** | DUF1249 (1) |
| **69** | DUF1501 (56) | **139** | DUF3455 (9) | **209** | DUF1326 (2) | **279** | DUF1223 (1) |
| **70** | DUF2715 (54) | **140** | DUF2846 (9) | **210** | DUF1028 (2) | **280** | DUF1043 (1) |

**Table S7.** Number of sequences from each type used in constructing the β-signal motif shown in Fig. 3C.

| | | | |
|---|---|---|---|
| **Prototypical** | 87,974 | **DUF1302** | 135 |
| **Fim/Usher** | 908 | **NfrA-like** | 71 |
| **LptD-like** | 1,596 | **PorT-like** | 106 |
| **LpxR-like** | 380 | **Hypothetical Protein Type 1** | 478 |
| **Tsx-like** | 103 | **Hypothetical Protein Type 2** | 76 |

**Dataset S1 (separate file).** FASTA file with the 121 sequences of the TMBB121 dataset.

**Dataset S2 (separate file).** FASTA file with the 1,000 sequences of the NONTMBB1K dataset.

**Dataset S3 (separate file).** FASTA file with the 1,929 sequences of the SS1929 dataset.

**Dataset S4 (separate file).** FASTA file with the 29,183 sequences of the TMBB29183 dataset.

**Dataset S5 (separate file).** FASTA file with the 651,874 sequences of the OMPDB-NR dataset.

**Software S1 (separate file).** IsItABarrel algorithm (Python script isitabarrel.py).
